# Storm in a bottle: An experimental investigation of how extreme precipitation impacts phytoplankton communities

**DOI:** 10.1101/2025.05.20.655176

**Authors:** Alex Barth, James L. Pinckney, Julie Krask, Erik Smith, Joshua Stone

**Affiliations:** University of Texas at Austin, Marine Science Institute, 750 Channel Drive, Port Aransas, Texas, USA; University of Texas at Austin, Statistics and Data Science; University of South Carolina, Biological Sciences, 700 Sumter St 401, Columbia, SC 29208; University of South Carolina, Belle W. Baruch Institute for Marine and Coastal Sciences, Georgetown, SC, 29442

## Abstract

Phytoplankton community composition in estuaries is tightly linked to freshwater input. While regular freshwater input typically delivers nutrients, fueling phytoplankton growth, the impact of extreme events is less certain. Several observational studies have documented increases in phytoplankton biomass following large precipitation events but cannot to adequately identify the mechanism driving this increase. This paper advances two hypotheses about what drives phytoplankton biomass change following extreme precipitation events. First, the Resident Response Hypothesis (RRH), which suggests local estuarine phytoplankton grow in response to favorable conditions. Alternatively, the Production Introduction Hypothesis (PIH) indicates that large rainfall events introduce new phytoplankton taxa to estuaries during run-off events associated with large storms.

These hypotheses were tested at North Inlet Estuary (South Carolina, USA) through a novel experimental design, which utilized multiple treatments to mimic the distinct impacts of large rainfall events on natural phytoplankton communities. Experimental samples were analyzed using photopigments and flow-through imaging microscopy. This allowed a wide assessment of phytoplankton community change both through measuring chlorophyll-a and biomass concentration. Ultimately, there was a strong increase in phytoplankton growth in the storm treatments, primarily identified by diatom pigment increases in response to run-off delivered nutrients, providing support for the RRH. However, biomass concentration analysis of select diatom taxa revealed the introduction of new diatom taxa in run-off water communities. This supports the PIH, suggesting that initial growth from nutrient additions may be due to small cell growth as well as introduction of new taxa.

## 1 Introduction

As the intersection of rivers and ocean, estuaries are notorious for being dynamic ecosystems with strong oscillations in local environmental conditions. Rainfall delivers terrestrial run-off and increases riverine and freshwater inputs to estuarine ecosystems. While increases in run-off and riverine inputs may lead to eutrophication at times (Pinckney et al. 2001), it is a standard mechanism of nutrient delivery to estuaries, which stimulates primary production in otherwise nutrient-limited systems (Pinckney et al. 1998; Cloern et al. 2014). Despite significant differences in characteristics (e.g., residence time, size, tides) between estuaries, many studies have identified a strong linkage between freshwater input and phytoplankton growth (reviewed by Cloern et al. 2014). Ultimately this production supports higher trophic levels, as evidenced by studies which link coastal and estuarine fisheries to freshwater flows (Meynecke et al. 2006; Gillson 2011; Broadley et al. 2022). The propagation of primary production to higher trophic levels in the marine environment can depend in large part on the community composition of those primary producers.

Thus, understanding phytoplankton community regimes remains an active research effort. The nutrient and hydrological conditions created during regular freshwater input events can drive the growth of many phytoplankton groups. Specifically, diatoms and similar large phytoplankton cells have been shown to grow quickly in high-nutrient conditions in temperate estuaries (Pinckney et al. 1999; Dorado et al. 2015; Cloern 2018). This empirical evidence largely aligns with classic understandings of phytoplankton community patterns (Margalef 1978). However, during intense rainfall events and severe storms, the phytoplankton community response can be markedly different than standard successional patterns (Thompson et al. 2023). Understanding the phytoplankton response to extreme rainfall events and storms is critical, as these events are becoming more common. Increases in storm frequency and intensity are projected as consequences of ocean warming (IPCC 2022), with specific increases in extreme, short-duration precipitation events (Westra et al. 2014). The Southeastern United States is an area of particular concern, where increases in Atlantic hurricane activity will impact coastal regions (Bender et al. 2010). While it is difficult to attribute any single storm event directly to climate change, there is strong empirical evidence for increases in storms and flooding in this region (Paerl et al. 2019). These intense storm events bring massive precipitation and flooding to estuarine ecosystems, driving change across multiple levels of biological communities. Phytoplankton in particular, are directly impacted by these massive precipitation events as they are immobile and subject to changes in water quality and river flow dynamics.

A growing number of studies have documented the unique response of estuarine phytoplankton following severe storm events across the Mid-Atlantic and Southern US. In the Chesapeake Bay Miller et al. (2006) used remote sensing and pigment analysis to identify an untimely bloom of diatoms following Hurricane Isabel. Comprehensive studies of Galveston Bay following Hurricane Harvey documented a clear shift in the local water chemical composition, indicating strong terrestrial influences (Steichen et al. 2020). While the phytoplankton community was initially displaced, there was a response of phytoplankton growth across a number of taxa (Steichen et al. 2020; Quigg et al. 2024). Notably, there was an introduction of freshwater taxa to the system, which progressively declined over time (Quigg et al. 2024). Similar observations of large flushing followed by increased phytoplankton blooms have been documented in a Florida estuary (Hagy et al. 2006). In a lower salinity system, Lake Pontchartrain experienced large phytoplankton blooms, primarily composed of diatoms, following the successive hurricanes Katrina and Rita, which reduced over several weeks (Pinckney et al. 2009). Longer-term observations in South Texas have identified prolonged periods of storm events result in a persistent increase in phytoplankton biomass, coinciding with a shift in the dominant phytoplankton community (Reyna et al. 2017; Douglas et al. 2023). These examples are just a few of the many observational studies that have documented the phytoplankton response in estuaries and coastal systems following extreme storm events. Generally, most of these cases exemplify some mixture of displacement, growth, and community composition change. Yet, the extent of these impacts is governed not only by local estuarine characteristics but also the unique attributes of individual storms (Wetz and Paerl 2008). Observational studies, while useful in their ability to document natural changes, are rarely able to disentangle the different mechanistic effects a storm has on the phytoplankton community. As extreme storms and their associated precipitation events increase, it is critical to develop a comprehensive understanding of phytoplankton community response to these major events.

In this paper, we present an experimental study to determine how natural estuarine phytoplankton communities are impacted by the different mechanisms of intense rainfall events. During an extreme rainfall event and the subsequent run-off and increased riverine flow, there are several concurrent mechanisms which impact the phytoplankton community. Observational studies often documented an increase in phytoplankton biomass, usually evidenced by chlorophyll-a, yet this could be due to different community changes among the phytoplankton. We advance two potential hypotheses to explain increases in phytoplankton biomass following severe rain events. First, the Resident Response Hypothesis (RRH): large rainfall events create favorable conditions for phytoplankton growth, and the preexisting estuarine, resident community grows in response. Multiple mechanisms, including a reduction in microzooplankton, could drive the RRH itself through grazing dilution salinity changes which favor specific taxa, or nutrient addition (the most common explanation in observational studies and phytoplankton growth paradigms). Alternative to the RRH, an observed increase in phytoplankton biomass may be driven by the introduction of new taxa - the Production Introduction Hypothesis (PIH). To test these hypotheses, an experiment was designed to mimic an extreme precipitation event, separating the distinct mechanisms acting on the phytoplankton community. We conducted this experiment in the North Inlet estuary in South Carolina, USA using natural communities of phytoplankton.

## 2 Methods

### 2.1 Study Location & Experimental Design

This study was conducted at the North Inlet-Winyah Bay National Estuarine Research Reserve (NIWB NERR) in August 2023 at Oyster Landing, North Inlet (33°21.1 N, 79°11.2 W). North Inlet is a salt marsh estuary with strong tidal cycles and short residence times (Allen et al., 2014). The estuary is a small system (32 km^2^) which drains a relatively pristine, small watershed composed of mixed forested wetland and upland bounded by *Spartina spp.* salt marsh. Regular freshwater input is small ( 10m^3^s^-1^) (Dame et al., 1991; Allen et al., 2014). The average salinity values typically oscillate around 30-32 during non-storm conditions. However, during periods of heavy rainfall, the salinity in the upper creek systems can reach near freshwater levels with sustained values around 10-15 ppt. Thus, it is an ideal location to investigate the impact of severe rainfall events in isolation as the confounding variables of anthropogenic run-off are primarily removed. While scattered showers are a common phenomenon, there have been several extreme rainfall events including notable hurricanes. Following these extreme rainfall events, salinity at Oyster Landing can plummet to mesohaline levels (<15) while the strong tidal signal drives mixing of freshwater inflow with the otherwise oceanic water masses.

To mimic an extreme rainfall event under experimental conditions, high tide (oceanic; salinity >28) water from Oyster Landing was collected before a light rainfall event. Then, flowing water from the nearest freshwater marsh (salinity ∼0.2; hereafter referred to as marsh water) was collected just before a drainage culvert where it would mix into higher salinity water (below drainage ditch was >10). Both seawater and freshwater samples were immediately siphoned through a 153µm mesh to gently remove any large zooplankton predators. To separate the different potential rainfall mechanisms, five experimental treatments were created using a 50%-50% mixture of: A complete control of 100% (no mix) sieved seawater (CTL), a mixture of sieved seawater and 0.2µm filtered seawater to test microzooplankton predator dilution (DIL), a mixture of sieved seawater and DI water to add salinity change (SAL), a mixture of sieved seawater and 0.2µm filtered marsh water to add inorganic nutrients from the marsh water (NUT), and finally a mixture of sieved seawater and sieved marsh-water to add any potential freshwater phytoplankton community (COM). These five treatments effectively act as a set of cascading controls where an additional mechanism of a storm is added in each successive treatment. If phytoplankton grew or declined in one treatment, but not in the prior ones, it would indicate that the added mechanism is driving the change. In detail, the DIL treatment tests if a relief of microzooplankton grazing pressure increases phytoplankton growth; the SAL treatment contains that grazing relief and tests if salinity change influences the phytoplankton community; the NUT treatment adds a test for whether inorganic nutrients increase phytoplankton growth; and finally, the COM treatment adds a test for the introduction of freshwater taxa. Thus, by comparing across treatments, we can identify the mechanisms of action driving change to the phytoplankton community, while still utilizing natural communities. All treatments were replicated five times at a total volume of 1 L for each replicate (mixtures were 500 mL of sieved seawater and 500 mL of treatment). Replicates were mixed and housed in acid-washed, clear PETG media bottles (Nalgene, ThermoFisher Scientific, Waltham, MA, USA. catalog number: 342020-1000). Then, the culture bottles were placed in large, hard-mesh boxes and suspended in the upper layer (top 1m) of water at Oyster Landing for a 48-hour in-situ incubation. This approach allowed for the culture bottles to be naturally mixed by tides, winds, and passing boat traffic. Additionally, the samples were all kept under natural and similar temperatures and lighting conditions. Before incubation, initial samples were taken from each treatment to process as a baseline condition following the same protocol as the final samples (see below for phytopigment and imaging protocols). Sample aliquots from each of the five treatment conditions (CTL, DIL, SAL, NUT, COM) were analyzed for inorganic nutrient concentrations on a SEAL AA3 Segmented Flow Analyzer (SEAL Analytical, Miluakee, WI, USA) (Supplemental Figure 1). Specifically, inorganic nitrogen species ammonium (NH_4_), nitrate + nitrite (NO_x_), and orthophosphate (PO_4_) were measured using the standard indophenol blue and nitrate reduciton by spongy cadmium methods, respectively, while orthophosphate was determined via the molybdenum blue method (Eaton et al., 2005). These measurements confirmed the experimental design - the sieved seawater had higher levels of ammonium (NH_4_) and orthophosphate (P0_4_) yet lower nitrate + nitrite levels (NO_x_), and the added marsh water treatments had high levels of nitrate. North Inlet is a generally nitrogen-limited system (Pinckney et al. 2020; Schlenker and Pinckney 2025), thus the final two treatments (NUT, COM) added a nutrient mechanism as desired. The DIL and DIW treatments did not have this nutrient enrichment. Following the 48-hour incubation, cultures were collected and immediately placed on ice in a dark cooler, then transported back to the lab for processing. Each experimental replicate was split for processing with high performance liquid chromatography (HPLC) to measure phytoplankton pigments or with a FlowCam imaging microscope. First, a well-mixed sample was taken from each experimental replicate and vacuum filtered onto 2.5 cm GF/F Whatman glass microfiber filters. For all replicates, the total volume for the HPLC sample was 300 mL except for the COM-treatment which was too concentrated and only 250 mL was taken to avoid clogging the filter. Filters were stored at -80°C. Then the remaining sample (700-750 mL) from each bottle was concentrated using a 20µm mesh sieve and suspended in 123mL of filtered water matching the salinity of the sample. These samples were then stored in amber bottles with a small amount (2mL) of Lugol’s solution added for preservation. Preserved samples were stored in the dark at room temperature for later imaging (within 3 months).

### 2.2 Phytoplankton Pigment Analysis

HPLC was used to determine the phytoplankton community across experimental conditions. This can provide measurements of chlorophyll-a as well as the accessory pigments attributable to different phytoplankton taxonomic groups (e.g., diatoms, dinoflagellates, etc.).

Chlorophylls and carotenoid photopigments were measured using high-performance liquid chromatography (HPLC). Sample filters were lyophilized for 24 hours at -50° C, placed in 90% acetone (1 ml), and extracted at -20°C for 18 to 20 hours. Extracts were filtered (0.45 µm, 250 µl) and injected into a Shimadzu 2050 HPLC system (Rainin Microsorb-MV, 0.46 x 10 cm, 3 µm and Vydac 201TP54, 0.46 x 25 cm, 5 µm columns). A nonlinear binary gradient of 80% methanol to 20% 0.50 M ammonium acetate and 80% methanol to 20% acetone was the mobile phase (Pinckney et al. 1996, 2001). Spectra and chromatograms (440 ± 4 nm) were identified by comparing retention times and absorption spectra with photopigment standards (DHI, Denmark). The relative concentrations of major algal groups based on the measured photopigment concentrations were estimated using ChemTax (v. 1.95) (Pinckney et al., 2001; Higgins et al., 2011). Total chlorophyll-a was partitioned into the concentrations of algal groups (e.g., diatoms, cyanobacteria, cryptophytes, etc.). The initial ratio matrix randomization procedure with 60 simulations was employed to minimize errors in algal group biomass resulting from inaccurate pigment ratio seed values (Higgins et al., 2011).

### 2.3 FlowCam Imaging

Generally, taxonomic identification and abundance estimation of phytoplankton communities is a time and labor-intensive process. However, high-throughput imaging microscopes can significantly alleviate this challenge by imaging a substantial volume of water (relative to what is feasible with light-microscopy via the Utermöhl method) in just a few minutes. For this experiment, we utilized a FlowCam 8000 (Yokogawa Fluid Imaging Technologies) with the following specifications for sample preservation and instrument configuration (Owen et al., 2022). Samples were preserved in Lugol’s solution as described above. Each sample was imaged in auto-image mode using a syringe pump with manufacturer default settings. Both the 10x and 4x objective to target different-sized particles. For the 10x objective, samples were pre-filtered using a 100µm mesh to reduce clogging of the flowcell. Then, because a 20µm mesh was used to concentrate samples on the initial processing, this was set as the minimum detection size (measured as area-based diameter, ABD) for the image segmentation in the VisualSpreadsheet software (version 6). For the 4x objective, samples were pre-filtered using a 150µm mesh (although this step was largely redundant given all treatments were initially filtered at 150µm). The minimum acquisition size was set to 20µm ABD to overlap with the 10x sample; however most of these images were filtered out at the identification stage (see below). To ensure that a sufficient number of particles were imaged in different size classes (at least 1500 individuals), multiple imaging runs per sample were conducted.

### 2.4 Image Classification & Taxonomy

While FlowCam has a native software for imaging processing and taxonomy in VisualSpreadsheet 6, we used the more popular plankton taxonomy application Ecotaxa (Picheral et al. 2017). All individual images, referred to as ROIs (regions of interest; note that many authors use the synonymous term “vignettes”), were exported from VisualSpreadsheet. Then to get the images and metadata compatible with the EcoTaxa ecosystem and match feature terms, the R package FlowProcess was used (https://github.com/TheAlexBarth/FlowProcess). EcoTaxa is a valuable tool for plankton classification because it offers the ability to easily add deep learning classification through feature extraction with an SCN, a process which can improve predictive performance in plankton classifiers (Irisson et al. 2022). EcoTaxa has several deep feature models that leverage the millions of other images hosted on the web application. While there are other FlowCam projects on Ecotaxa, and an associated deep feature extractor, most users following this approach do not utilize VisualSpreadsheet for image segmentation but rather ZooProccess (Gorsky et al. 2010). As a result, the ROIs are different (ZooProcess provides only black-and-white while FlowCam-VisualSpreadsheet is colored). For this reason, we used the “planktoscope_2022-09” deep feature model. While the camera is slightly different, most PlanktoScope (Pollina et al. 2022) users utilize MorphoCut for segmentation, which preserves the color features. Plankton classification was done separately for the 4x objective and the 10x objective. For the 4x objective, only ROIs larger than 70µm ABD were considered as those smaller than this were generally not identifiable to a useful level. In total, this resulted in 94,992 and 1,084,152 ROIs for the 4x and 10x objectives respectively. Since it would not be feasible to manually sort all these images, we established a reliable method to utilize the classifier for predicting different phytoplankton taxa, as other studies have done with imaging microscopy (Quigg et al. 2024). For both magnification levels, a custom learning set was developed based on 10% of the total images. Using the learning set, the remainder of the data were predicted. For this, 10% of the predicted data was randomly selected for manual validation to assess classifier performance. To be conservative in testing the experimental hypotheses, only taxa which consistently were able to be identified through accurate positive predictions were utilized (see Supplemental Information 2). Following this critera, only 7 taxonomic or morphological groups of diatoms were used in this study; *Rhizosolenia spp.*, *Skeletonema spp*., *Entomoneis sp.*, *Cylindrotheca sp*., Unindentified Centrics (likely mostly *Thalassiosira spp.* individuals), *Coscinodiscus spp.*, and *Corethron sp* (Figure 1). Notably diatoms made up the vast majority of the learning and validation sets, thus by focusing on diatoms we are likely capturing the bulk of the phytoplankton community change in this size fraction. However, because we are relying on positive predicted data, combined with biases of Lugol’s preservation (Zarauz and Irigoien 2008), these results should not be used for external estimation of total change (e.g., growth rate calculations) but rather are only useful for assessing relative change within the context of this experimental study.

Data were exported from EcoTaxa for analyses. For each ROI of an individual diatom, biovolume (µm^3^ Cell^-1^) was estimated using ABD, which has been shown to perform similarly to more complex shape-based volume estimation methods (Jakobsen and Carstensen 2011; Álvarez et al. 2014; Hrycik et al. 2019). Then biovolume to carbon-biomass pgC Cell^-1^ were calculated following standard log-log equations (Menden-Deuer and Lessard 2000). Different equations were used based on cell size, greater than or less than 3000µm^3^. Notably, although some *Skeletonema spp.* chains were larger than 3000µm^3^ all were treated as small since the biovolume of individual cells is small regardless of the total chain size. Then within each experimental replicate, the carbon biomass concentration (pgC mL^-1^) was calculated for each diatom group based on total carbon biomass divided by the volume of water processed.

**Figure 1:**
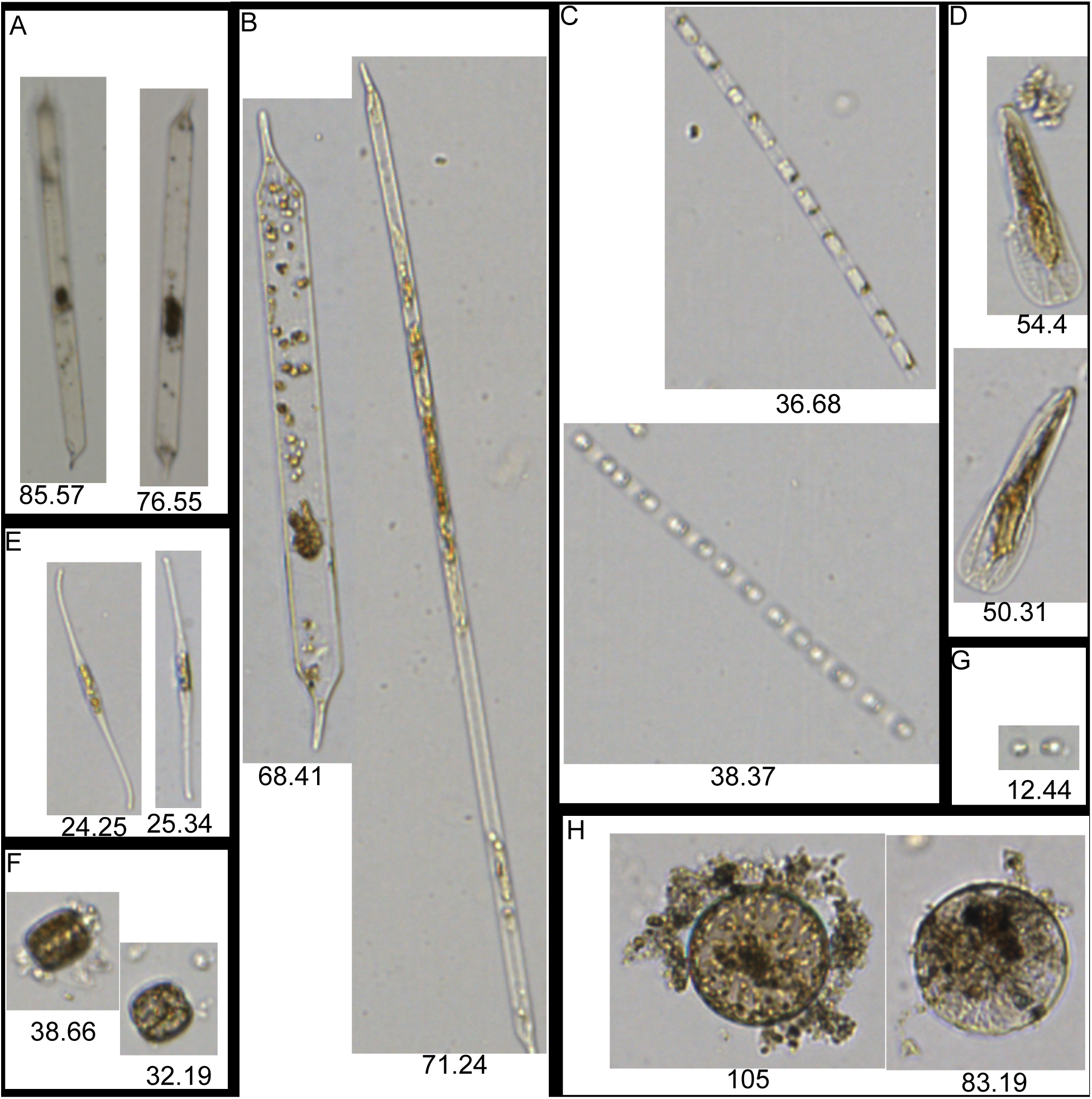
Example images of Diatoms imaged by FlowCam and successful predicted through EcoTaxa classification. Imaged with 4x magnification (A) Rhizosolenia; Imaged with 10x magnification (B) Rhizosolenia, (C) Skeletonema, (D) Entomoneis, (E) Cylindrotheca, (F) unclassified Centric diatom, (G) Corethon, (H) Coscinodiscus

### 2.5 Statistical analysis

For both HPLC measurements (chlorophyll-a and ChemTax derived estimates) and biomass concentration analysis, two metrics were assessed: (1) final values in each replicate after the 48-hour incubation and (2) the change in values for each replicate relative to the initial state. Both metrics are necessary for thoroughly evaluating the hypotheses. Assessing the change from the initial state will inform growth (or loss), allowing the test of the RRH. By assessing the final values, it is possible to test for successful introduction of new phytoplankton communities (PIH). While it is possible to assess the introduction from measurements of the initial state, looking at the end values informs a successful introduction, in which new community members survived the addition to a mesohaline environment for at least 2 days.

For all data types (HPLC or FlowCam), the same statistical model was used to estimate the mean effect of each treatment. Observations of the final conditions (*y_i,g_*) for the ith replicate of each treatment, g, were modeled as:

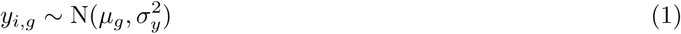

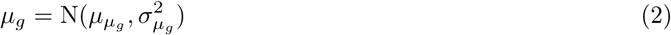

This linear model is conceptually similar to an ANOVA, where the data model (Equation 1) assumes all replicate observations come from a treatment-specific mean effect (*µ_g_*) and homogeneous variance in observations across treatments. Then, using a Bayesian approach, inference can be made on the mean treatment effects. A weakly informative prior was set for each mean effect (Equation 2). The prior mean and variance was the same for each treatment (g), just adjusted to be on the scale of the data (chlorophyll-a or carbon biomass). A similar model was constructed for inferring the treatment mean difference between final and initial conditions:

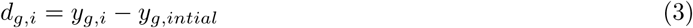

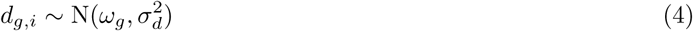

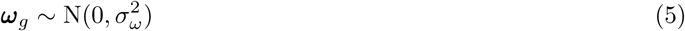

Where the observed difference (*d_g,i_*) was calculated (Equation 3), then modeled from a normal distribution with a treatment-specific mean difference effect (Equation 4). Again, weakly informative priors were set with prior treatment variance on the scale of the data, although for all analyses the prior mean was 0, implying a weak prior assumption of no-effect. All analyses were conducted in R (version 4.4.0) and posterior estimates were calculated using the Hamiltonian Monte-Carlo software Stan (version 2.32.2). All code and related data products are publicly available (INSERT GITLINK ON PUBLICATION).

## 3 Results

### 3.1 Chlorophyll-a and Phytoplankton Pigments

The total chlorophyll-a, measured by HPLC, showed clear patterns across treatments. The final mean chlorophyll-a (Figure 2a) and the mean chlorophyll-a change (Figure 2b) were low for all non-marsh water treatments. Inference on the mean effect showed no clear difference between the control, dilution, and salinity change treatments (all 95% credible intervals overlap, Figure 2c,d). The control treatment had an evident decline in total chlorophyll-a relative to initial conditions. Alternatively, there was higher chlorophyll-a signal in the two marsh water treatments. The nutrient treatment had a final mean chlorophyll-a concentration of 10.20µg L^-1^ (95% Cred: 8.35, 11.76) while the composition treatment had a final mean chlorophyll-a concentration of 11.02µg L^-1^ (95% Cred: 9.08, 12.55). Both these marsh-water treatments had similarly high posterior estimates of the mean chlorophyll-a change (Figure 2d; Supplemental Information 3). However, for either analysis of total chlorophyll-a, there was no clear difference between nutrient treatment or the composition treatment (Figure 2, Supplemental Information 3).

Using the results from the ChemTax, it is possible to partition the total chlorophyll-a across broad phytoplankton taxonomic groups. Across all treatments, diatoms had a mean final chlorophyll-a concentration of 5.05 µg L^-1^ (2.98 standard deviation), contributing approximately two-thirds to three-quarters of the total community chlorophyll-a signal (Figure 3). The following most prevalent taxa were chlorophytes, cyanobacteria, and cryptophytes at an average of 16.3%, 5.41%, and 3.61% of the total chlorophyll-a signal across treatments. Dinoflagellates, prasinophytes, haptophytes, and euglenophytes each contributed less than 1% across treatments.

The posterior estimates of the final mean and mean change in chlorophyll-a for both diatoms and chlorophytes primarily aligned with the pattern of the total chlorophyll-a analysis (Figure 4a-d). This pattern for both groups showed no clear difference across the first three treatments (control, dilution, and salinity) for the final chlorophyll-a concentration. Additionally, none of the 95% credible intervals for the estimate of the mean change in chlorophyll-a suggested any growth in these first three treatments (Figure 4b,d; Supplemental Information 3). Alternatively, the final two marsh-water treatments (nutrient and composition) had significantly higher chlorophyll-a levels, yet not clearly different from one another (Figure 4a-d). The cyanobacteria and cryptophytes, which were relatively less abundant yet still prevalent, had a different pattern. cyanobacteria’s final mean chlorophyll-a was higher in the nutrient treatment relative to the non-marsh water treatments, and even higher in the composition treatment (95% posterior credible intervals non-overlapping between nutrient and composition treatments; Figure 4e; Supplemental Information 3). Furthermore, the cyanobacteria chlorophyll-a change analysis (Figure 4f), suggested no growth in the control or dilution treatments but progressively higher growth across the salinity, nutrient, and composition treatments (Supplemental Information 3). The cryptophytes similarly had a higher final mean chlorophyll-a estimates between the composition, nutrient, and three non-marsh water treatments. However the suggested growth by chlorophyll-a change for cryptophytes was not across treatments (Figure 4g-h; Supplemental Information 3).

**Figure 2:**
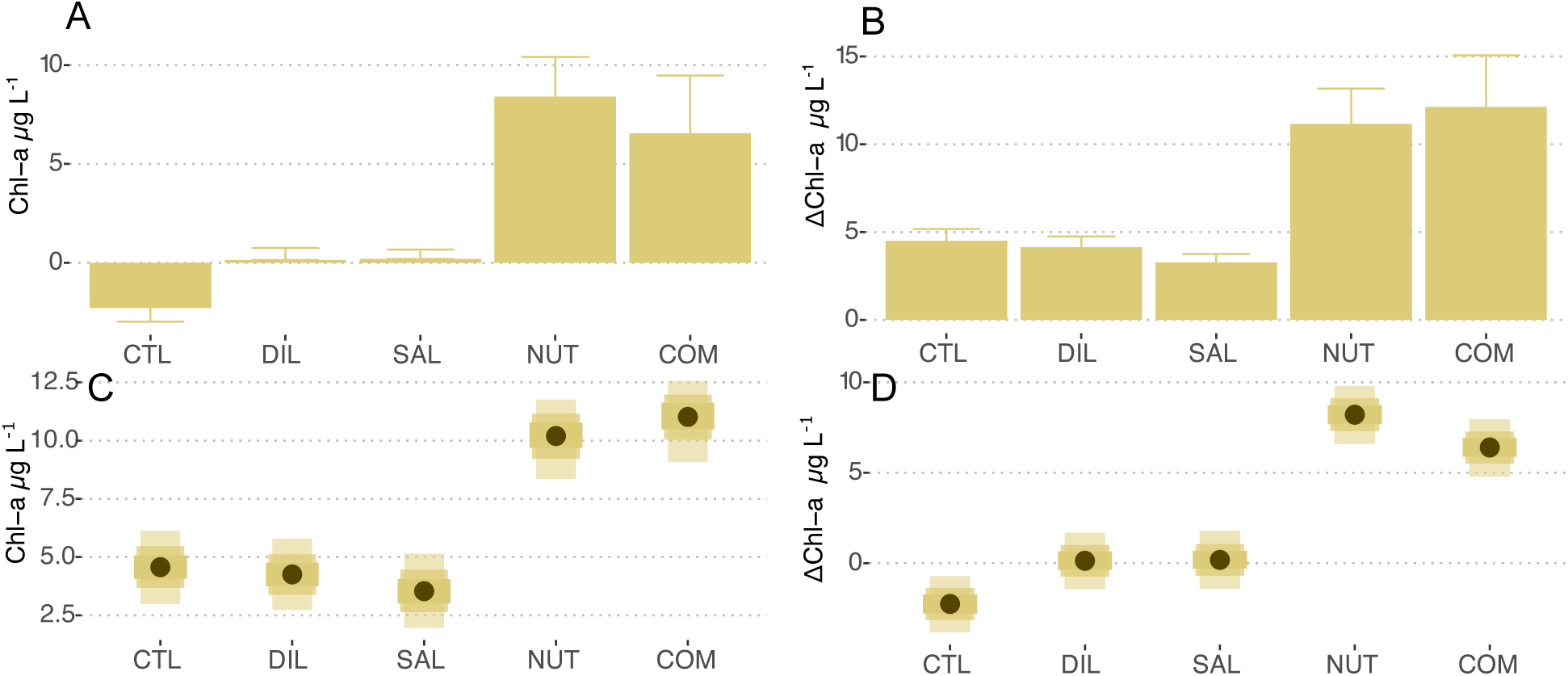
Total chlorophyll-a measurements through HPLC. (A) Mean final values between treatments with standard deviation shown. (B) Mean difference in each treatment relative to initial concentration with standard deviation. (C) Posterior credible intervals and posterior mean estimates of the treatment final concentration mean. (D) Posterior credible intervals and posterior mean estimates of the treatment mean difference. Shaded credible intervals in C,D, are at 50%, 75% and 95% equal-tailed density estimates with points representing the posterior mean.

**Figure 3:**
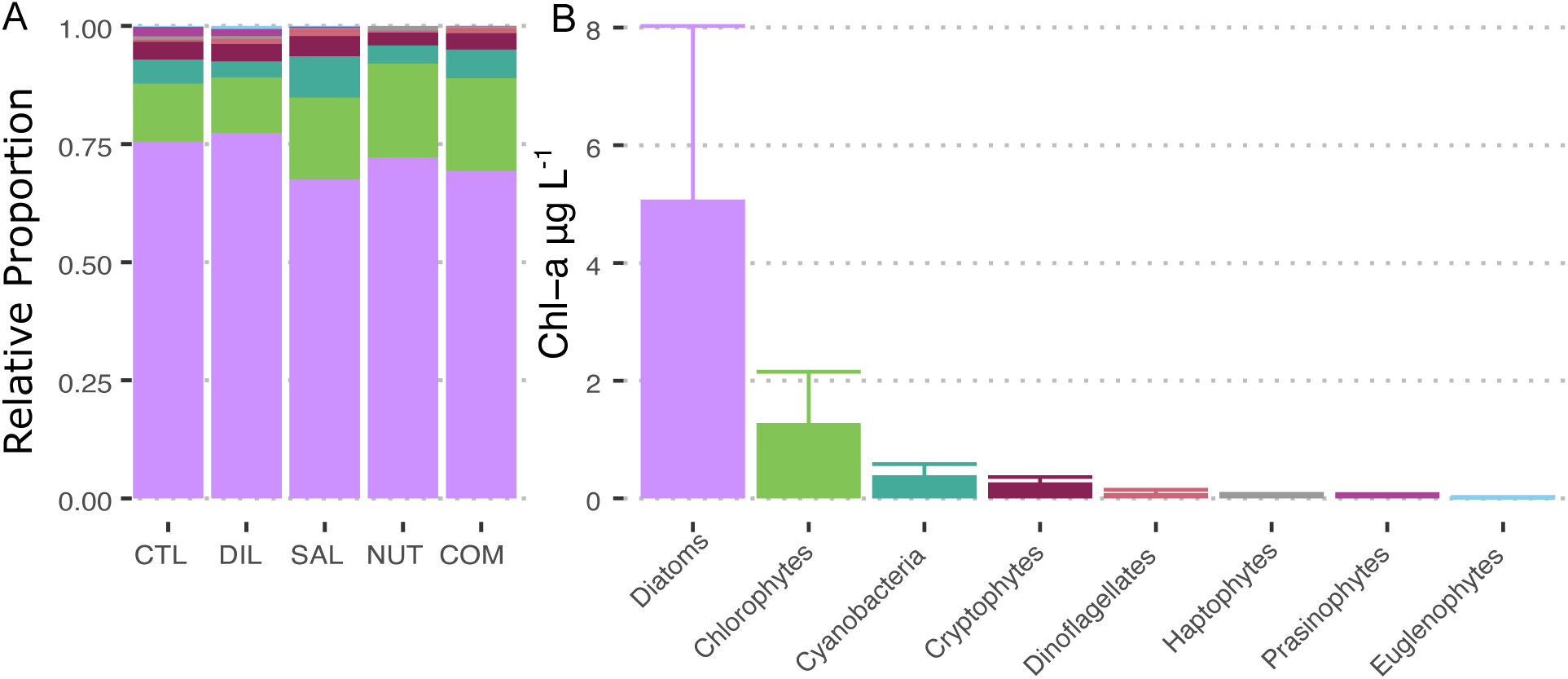
(A) Relative abundance of major phytoplankton groups’ contribution to total chlorophyll-a as measured by ChemTax in final concentration measurements between treatments. (B) Mean chlorophyll-a attributable to different taxonomic groups across all treatments shown with standard deviation. Color panel of (A) corresponds to taxonomic groups shown in (B).

### 3.2 Carbon Biomass Concentration

Only seven diatom taxa which were reliably identified by FlowCam imaging and EcoTaxa prediction were investigated (*Rhizosolenia spp.*, *Skeletonema spp*., *Entomoneis sp.*, *Cylindrotheca sp*., Unindentified Centrics (solitary morphological group), *Coscinodiscus spp.*, and *Corethron sp*). Of these groups, *Rhizosolenia spp.* was generally the most prevalent taxa across treatments, except the composition treatment (Figure 5a) with a mean final biomass concentration of 12.9 × 10^7^ pg C mL^-1^ (Figure 5b). Following in relative abundance to a lesser extent were *Coscinodiscus spp.* and *Skeletonema spp.*. *Cylindrotheca spp.* and *Entomoneis sp.* were both prevalent only in the composition treatment (Figure 5a).

**Figure 4:**
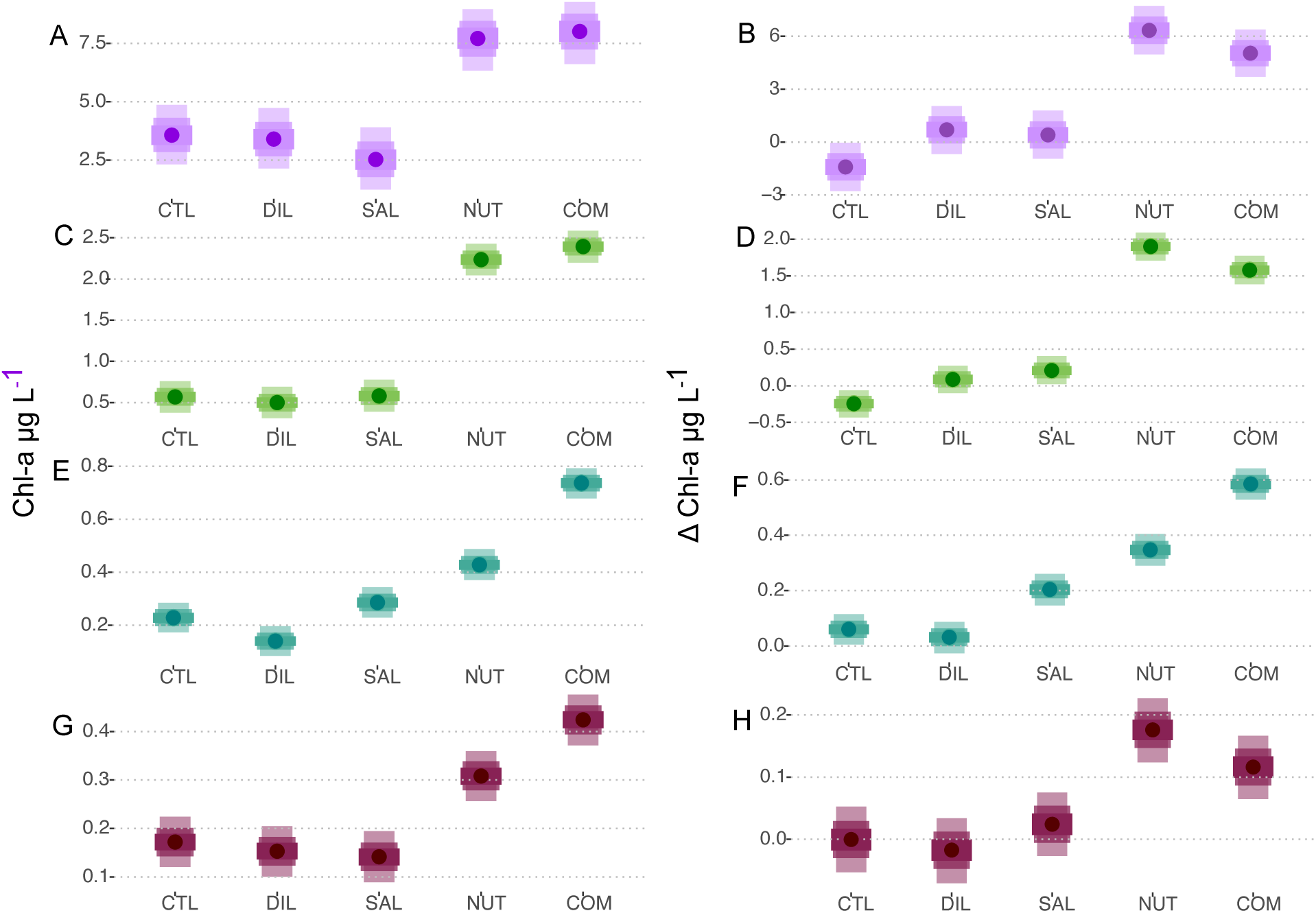
Posterior estimates of treatment effects estimated by ChemTax for the five most relatively abundant taxa (A,B) diatoms, (C,D) chlorophytes, (E,F) cyanobacteria, (G,H) cyrptophytes. Left panels (A,C,E,G) show the final mean chlorophyll-a concentration, while right panels (B,D,F, H) show a difference in chlorophyll-a concentration from the initial. Shaded credible intervals in C,D, are at 50%, 75% and 95% equal-tailed density estimates with points representing the posterior mean.

Statistical analysis of mean final and change in the biomass concentration across treatment had different results across taxa. *Rhizosolenia spp.*, had a slight trend of final concentration being higher in the non-freshwater treatments (control and dilution), however the 95% credible intervals overlapped for most treatments (Figure 6a; Supplemental Information 3) and there was no evidence of *Rhizosolenia spp.* growth across any of the treatments (Figure 6b). A similar pattern was observed in *Coscinodiscus spp.* and *Skeletonema spp* (Figure 6). Opposite to the bulk of the investigated diatom biomass, *Cylindrotheca spp.* and *Entomoneis sp.* displayed clearly higher final mean carbon biomass concentration in the final composition treatment. However, the growth of these two taxa in the composition treatment, although clearly higher than 0, was not clearly different than other treatments (Figure 6e-h). The less abundant taxa, Unidentified Centrics and *Corethron sp.* were not different across treatments (Supplemental Information 3).

**Figure 5:**
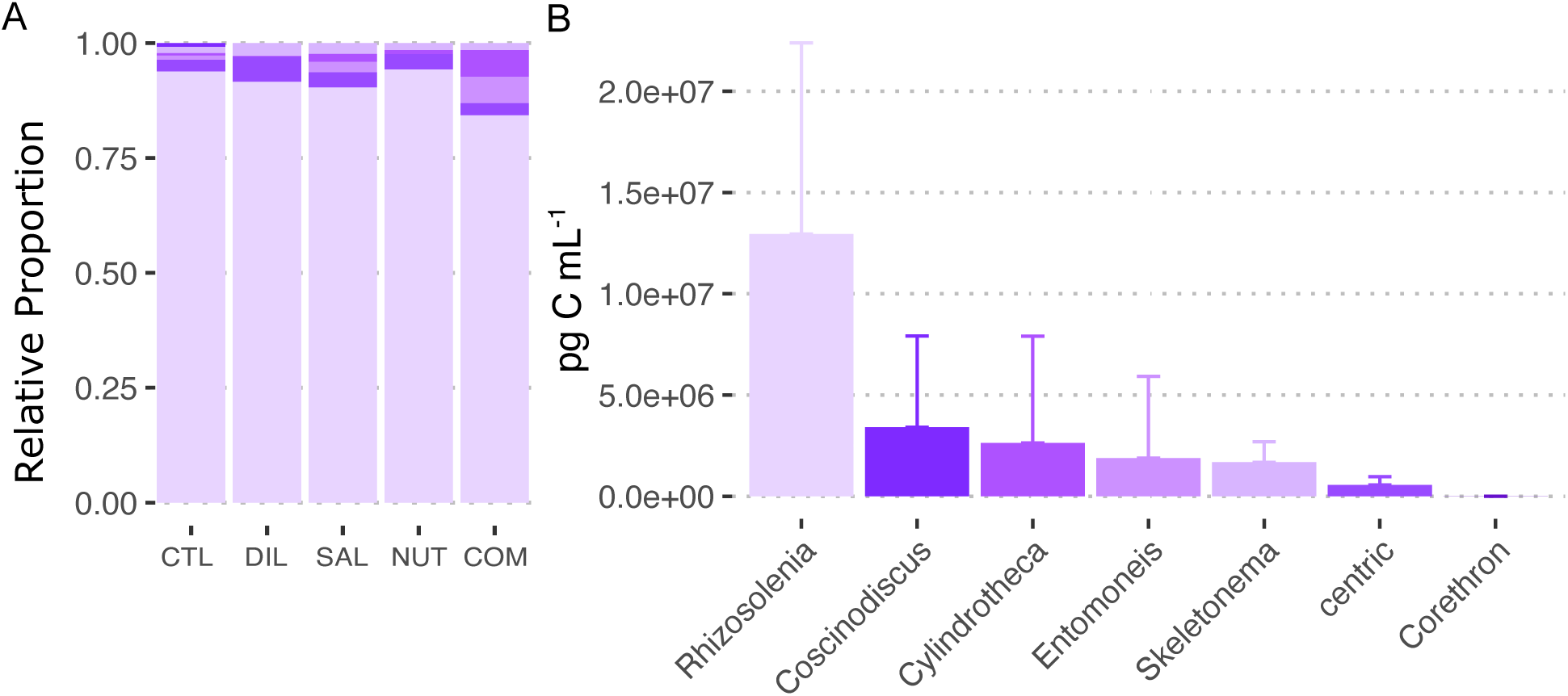
(A) Relative abundance of different diatom groups’ contribution to total biomass concentration across final values in experimental treatments. (B) Mean biomass concentration attributable to different diatom groups across all treatments shown with standard deviation. Color panel of (A) corresponds to taxonomic groups shown in (B).

**Figure 6:**
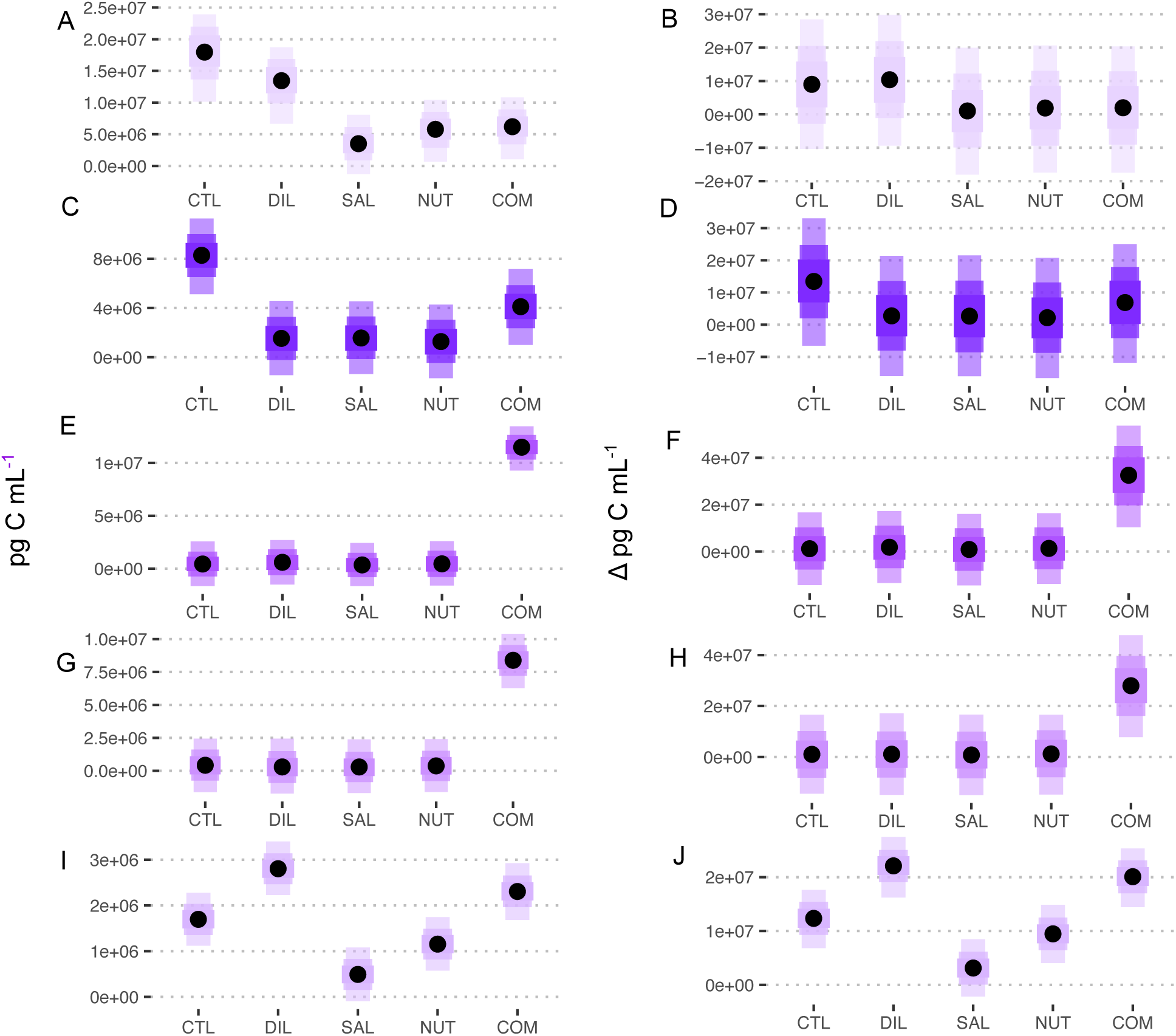
Posterior estimates of treatment effects for the five most relatively abundant diatom groups (A,B) Rhizosolenia spp., (C,D) Coscinodiscus spp. (E,F) Cylindrotheca sp., (G,H) Entomoneis sp and (I,J) Skeletonema sp.. Left panels (A,C,E,G,I) show the final mean biomass concentration, while right panels (B,D,F,H,J) show a difference in biomass concentration from the initial. Shaded credible intervals in C,D, are at 50%, 75% and 95% equal-tailed density estimates with points representing the posterior mean.

## 4 Discussion

This experiment effectively provided the opportunity to delimit the different mechanisms of an intense rainfall event to study the phytoplankton community response. As predicted by many observational studies, as the treatments progressively added the mechanisms of a storm, there was an observed increase in phytoplankton biomass. However, the point at which (and thereby the causal mechanism) this growth occurred was consistently clear across different analyses. We proposed two alternative hypotheses: the Resident Response Hypothesis (RRH) and the Production Introduction Hypothesis (PIH) to explain an observed increase in phytoplankton biomass following intense rainfall events. Chlorophyll-a and pigment analyses provided significant support for the RRH and minor support for the PIH. Alternatively, FlowCam-based image analysis only provided some support for the PIH. It should be noted that these hypotheses are not mutually exclusive, so the mixed evidence provided from this experiment is not inherently contradictory. Here, a broader context provides a clearer picture of how this experiment advances both hypotheses:

### 4.1 Resident Response Hypothesis

The analysis of the total chlorophyll-a signal largely supported the RRH through the mechanism of nutrient addition. The increase in chlorophyll-a occurred at the nutrient treatment level, but was not higher in the composition treatment. This suggests that the new nutrients (primarily nitrate, Supplemental Information 1) stimulated growth. This growth was evident among diatoms, which are the primary contributors to the total chlorophyll-a signal in this study. The basic paradigm that the introduction of nitrogen-rich freshwater into estuaries stimulates phytoplankton growth is a simple concept supported by a large number of studies (Margalef 1978; Pinckney et al. 1998; Cloern et al. 2014; Cloern 2018).

However, the evidence for this nutrient-introduction RRH gets cloudy when considering the full context of this experimental study. First, prior work in the US Southeastern estuaries noted the ChemTax algorithm does not always accurately predict phytoplankton biomass (Lewitus et al. 2005). Wetz and Paerl (2008) noted Diatoms and Chlorophytes were not well identified by ChemTax in a North Carolina estuary. Coincidentally, these were identified by our ChemTax analysis as the two most relatively abundant groups. Yet, even if ChemTax did not accureately represent the contribution of these groups, it doesn’t immediately nullify the observed support for the RRH. The total chlorophyll-a patterns, which support the nutrient-introduction mechanism of the RRH, are accurate measurements. Additionally, the FlowCam, while not able to measure smaller cells, supported the observation that diatoms were an extremely prevalent member of the phytoplankton community across all experimental treatments. However, none of the investigated diatoms in our analysis of the FlowCam data reflected pattern of nutrient-induced growth suggested by the analysis of chlorophyll-a. There are a few potential explanations for this discrepancy.

One consideration is that the observed increase in diatom pigments, and by extension, of ChemTax analyses, the contribution to the total community biomass was just an increase in pigments but not yet biomass. Even in the 0.2µm filtered marsh-water used in the nutrient treatment, there may have been an increase in color dissolved organic matter (CDOM) which would have shaded those replicates. Through photoacclimation, many diatoms are capable of adjusting their chloroplast and pigment concentration in response to low light conditions (Anning et al. 2000; Ghobara et al. 2019). The most prevalent diatom, of those investigated, was *Rhizosolenia spp.* which is a very large-bodied cell. Although some species of *Rhizosolenia spp.* have been documented to divide multiple times per day (Yoshimatsu et al. 2020), it can be much slower (Villareal 1990). Potentially, in the marsh-water treatments (NUT and COM), where the pigment analysis suggested an increase in Diatom biomass, there were large cells, like *Rhizosolenia spp.* that increased pigments but not biomass. Although this is a feasible explanation, it is not likely. All replicates were cultured in situ at Oyster Landing, which already has relatively high CDOM levels, likely creating a similar photic environment. Additionally, if shading by CDOM at the nutrient treatment level drove pigment increases, the same logic would suggest the additional shading by particulates in the composition treatment would further drive photoacclimation. Yet this was not observed. Finally, because these bottles were suspended in the upper layer of the water column, they received more sunlight than natural phytoplankton that are mixed throughout the water column (and experiencing periods of darkness), so the culture bottles may not even have an increase in overall shading.

A more plausible explanation to reconcile the conflicting evidence for the RRH is simply that the chlorophyll-a response was by cells not identified in our FlowCam imaging to EcoTaxa prediction pipeline. For instance, *Chaetoceros spp.* is prevalent in this system, and many species in this genus can multiple two or more times per day in ideal growth conditions (McGinnis et al. 1997; Sánchez-Saavedra and Voltolina 2006). While various *Chaetoceros spp.* were identified in initial FlowCam learning set development, their chains were often broken in preservation/storage, and their spines caused them to be aggregated with other particles, prohibiting their inclusion in a reliable predictive set. Another possibility is that in the favorable nutrient conditions created in the experimental marsh-water treatments, diatom resting spores were activated into vegetative states, a process which contributes to bloom formation following storm events in tropical estuaries (Patil and Anil 2008). Presumably, the newly-formed vegetative cells would be small and not readily identified in the imaging prediction pipeline employed in the present experiment. Regardless of the exact species or life stage, which contributed to the observed diatom growth, it is highly likely the observed pattern in our experiment was real biomass growth. In the weeks following Hurricane Harvey (2017), studies in Gavleston Bay noted large amounts of fucoxanthin (diagnostic pigment for diatoms) (Quigg et al. 2024). While an imaging analysis did not observe the most significant increase in diatom cell density until nearly 4 weeks following the initial storm (Quigg et al. 2024), 18S rRNA analysis documented a sizable increase in the relative abundance of Diatoms within two weeks (Steichen et al. 2020). While genetic-based estimates of relative abundance have unique biases and may not readily correlate with actual cell density, diatom growth might precede the point at which large cells are observable through flow-through microscopy.

### 4.2 Production Introduction Hypothesis

Both methods of measuring phytoplankton biomass provided some support for the PIH. First, the HPLC and ChemTax results demonstrated a clear process by which cyanobacteria were introduced and grew under experimental conditions. The salinity treatment, where just DI water was added, saw a higher growth in cyanobacteria (Figure 4f). The initial cyanobacteria in the salinity treatment originated from the high-salinity Oyster Landing water, which would suggest it would be salt-adapted (Moisander et al. 2002), however, many cyanobacteria strains thrive in lower salinity environments (Paerl and Otten 2013) so even if the initial community was salt-adapted, the reduction in salinity stress may have promoted favorable growth conditions. Then, in the nutrient treatments, the favorable salinity and nutrient-rich conditions likely furthered the ability for these cells to grow. Finally, the composition treatment clearly introduced additional cyanobacteria. Cyanobacteria were generally not identified by the FlowCam analysis, which is not surprising as these taxa are generally much smaller than diatoms and may not be identifiable at 100x magnification. Anecdotally, in the learning set development only two colonies of *Merismopedium* were identified, both of which belonged to the composition treatments. This colonial cyanobacteria, along with others, has been commonly identified in estuarine water post-storm events (Claflin et al. 2024; Quigg et al. 2024).

The FlowCam/EcoTaxa analysis also provided support for the PIH with two of the investigated Diatoms. *Entomoneis sp*. is a very distinguishable cell with a unique morphology. Several papers in the Eastern Hemisphere have described observations of *Entomoneis spp.* across marine, brackish, and freshwater environments (Liu et al. 2018; Mejdandžić et al. 2018). While research in Galveston Bay has identified *Entomoneis spp*, with particular increases following storms in 2017 (Quigg et al. 2024), it is generally not widely described in observational phytoplankton research. Many species of *Entomoneis spp.* are benthic and common in tidal flats (Jauffrais et al. 2016). It is likely that during the marsh-water collection for this experiment, the flowing water included suspended microphytobenthos, such as *Entomoneis spp.*, which were introduced and then grew in the composition treatments. A similar mechanism could be the case for *Cylindrotheca sp.*, which although prevalent in the pelagic is also a common member of benthic diatom assemblages. Thus, heavy rainfall events could not only introduce freshwater taxa to estuaries, but suspend benthic ones - suggesting a complex linkage across different compartments of the marine ecosystem.

### 4.3 Considerations and Conclusions

This study provided a robust experimental analysis to identify the unique impact of different mechanisms of severe rain events on local estuarine phytoplankton. We provide experimental evidence that new nitrate introduced from freshwater stimulates phytoplankton growth (RRH) during intense rainfall conditions, as suggested by many observational studies. The experiment also demonstrated that this growth was not due to microzooplankton grazing relief nor salinity change. Also, it is clear that freshwater or potentially benthic taxa can be introduced during storm conditions (PIH). Still, the context of this experiment should be considered before extrapolating these findings. On a local scale, residence time in North Inlet is very low. By design, this experiment replicated a residence time 48 hours, so results may be more reflective of estuaries with longer residence times. Additionally, the experimental treatments were conducted with natural phytoplankton communities. While this approach facilitates a realistic investigation of the whole-phytoplankton community response, results will be sensitive to the local starting conditions. For instance if a different seed community of taxa from either the control or full community treatment were prevalent, growth dynamic are likely to change. Another artifact of the experimental design is that the conditions, by necessity mimicked, a strong mixing of water. However, horizontal stratification may occur where the freshwater input following storms does not immediately mix with the higher salinity water masses in estuaries (Cloern et al. 2017). Lastly, during an intense rainfall event, the added water may dilute freshwater nutrients to be in lower concentration than what was recreated in experimental treatments <Was Gabby working on this?>. However, there is also the potential of atmospheric deposition of nutrients directly through rainfall mechanisms which may increase nitrogen addition (Paerl et al. 1990). Ultimately, these considerations highlight the importance of understanding local-scale processes in estuaries, which can be highly variable across different systems. Nonetheless, we present here a clear pattern through which severe rainfall events impact estuaries across a variety of mechanisms. We encourage researchers to expand upon this experimental design and replicate in different systems. Using a robust experimental design across systems combined with adaptive sampling surrounding storms (Wetz and Paerl 2008; Quigg et al. 2024) will significantly contribute to an improved understanding of how storm events impact the base of estuarine ecosystems.

## Acknowledgments

We appreciate the field, lab, and housing support provided by the Baruch Marine Field lab and the North Inlet-Winyah Bay National Estuarine Research Reserve. Specifically, we are grateful to B. Pfirrmann for experimental design help, N. Mallick for field assistance, and G. Madgett for HPLC processing in the Estuarine Ecology Lab. This work was funded in part by the University of South Carolina’s SPARC Graduate Research Grant Program and the Slocum-Lunz Foundation. Use of the FlowCam was made possible by Yokogawa Fluid Imaging Technologies, Inc. through the FlowCam Aquatic Research Equipment Education Grant for Graduate Students.

