## Supporting Information 1 for "Storm in a bottle: An experimental investigation of how extreme precipitation impacts phytoplankton communities"

Alex Barth

April 2025

Table 1: Nutrient values in initial treatment conditions. Note these are the treatment addition to control water, not the mixture which was cultured in each experimental chamber.

| Treatment | NO23 (uM) | NO2 (uM) | NH4 (uM) | PO4 (uM) |
|---|---|---|---|---|
| CTL | 0.581 | 0.366 | 8.211 | 1.103 |
| DIL | 0.255 | 0.015 | 0.183 | 0.014 |
| SAL | 0.722 | 0.185 | 1.115 | 0.801 |
| NUT | 3.658 | 0.871 | 5.948 | 0.251 |
| COM | 3.824 | 1.038 | 5.907 | 0.396 |
