## Supporting Information 2 for "Storm in a bottle: An experimental investigation of how extreme precipitation impacts phytoplankton communities"

### Taxonomy Supplement

#### Learning set development

Classficiation of FlowCam ROIs were done using Ecotaxa. In general all ROIs contained single particles (cells, detritus, or unidentifiable material). In cases where multiple cells were clustered these were classified as individuals since biovolume is the ultimate metric calculated. In cases where multiple organisms of different types were clustered, these were caterogirzed as "multiple other"

In total, the flowcam data processing yeilded 1,084,152 and 1,152,717 ROIs for the 10x and 4x magnifications respectively. To reduce this massive amount of images, we filtered the images to only include ROIs larger than 70$\mu$m for the 4x magnification as these largely were too small to identify and overlaped with the images collected by the 10x lens. On these filtered image sets, we utilized the deep-learning classificaiton from the popular Ecotaxa platform. For each magnificaiton setting, a custom learning set was developed based on 10% of the filtered images. This resulted in learning sets of just over 108k images in the 10x magnification and approximately 10k images in the filtered 4x magnification.

Generally, learning set criteria was based on the taxonomic references below and with some additional comments:

##### Phytoplankton:

Generally, diatoms were identifiable to more specific levels. However, there were a few instances of consistently unidenfiable cells. In cases of unidentifiable diatoms, they were grouped to a morphological class (e.g., centric or penate diatom). *Coscinodiscus spp.* were a well predicted group in both magnification levels. However, these often had a large amount of detritus surrounding individual cells. As a result, the biovolume estimates for this group are likely overinflated, but consistent for comparison across the experimental study. *Chaetaceros spp.* chains were a common group however, due to their spines, they were typically aggregated in a large mass of detritus, making consistent prediction challenging and were thus excluded from the analysis.

Other phytoplankton taxa were either less common or unidentifiable after Lugol's preservation. Some dinoflagellates were identified as thecated or naked but in total, they comprised less than 1/1000th of the learning set. The only reliably identified cyanobacteria was *Merismopedia*, a large filamentous mat. However, only three colonies were observed across both lenses.

##### Microzooplankton:

At the 4x magnification, a number of microzooplankton were identifable. Predominately loricated cells from *Choreotrichia.* These were grouped as *Eutintinnus spp.* and *Tintinnida.* Notably, many of these cells were empty lorica, however this is not unsurprising and many microzooplankotn do not preserve well in Lugol's. Small copepods, often late stage juveniles and their nauplii were also identified, but they were generally small in number.

##### Detritus:

To improve classifier performance, a number of detritus categories were created. However, these categories were not used in the learning set evaluation since they are not of ecological interest in this study.

#### Learning set evaluation:

Evaluation of classifier preformance used a randomly selected testing set comprised of 10% of the remaining unclassified images. The testing set was evaluated for True Positivies (TP) and False Positives (FP).

$$PPV = \frac{TP}{TP + FP}$$

Precision was calculated as the Positive Predictive Value (PPV, Equation 1) to quantify the preformance of the classifier. Each predicted image has a confidence score from the classifier. PPV was calculated across a range of confidence scores to identify a threshold at which predictions were reliable for distict categories.

##### 10x Precision:

Some categories performed strongly while others were disproportionaly weak (Figure 1). For instance, *Skeletonema sp.* and *Rhizosolenia spp.* were fairly precise "Centric Chains" had a very large number of FPs with high confidence scores. The poor preformance of a broad category like “Centric chain” is not surprising as this group contained a variety of cells which were too broken or unfocused to identify.

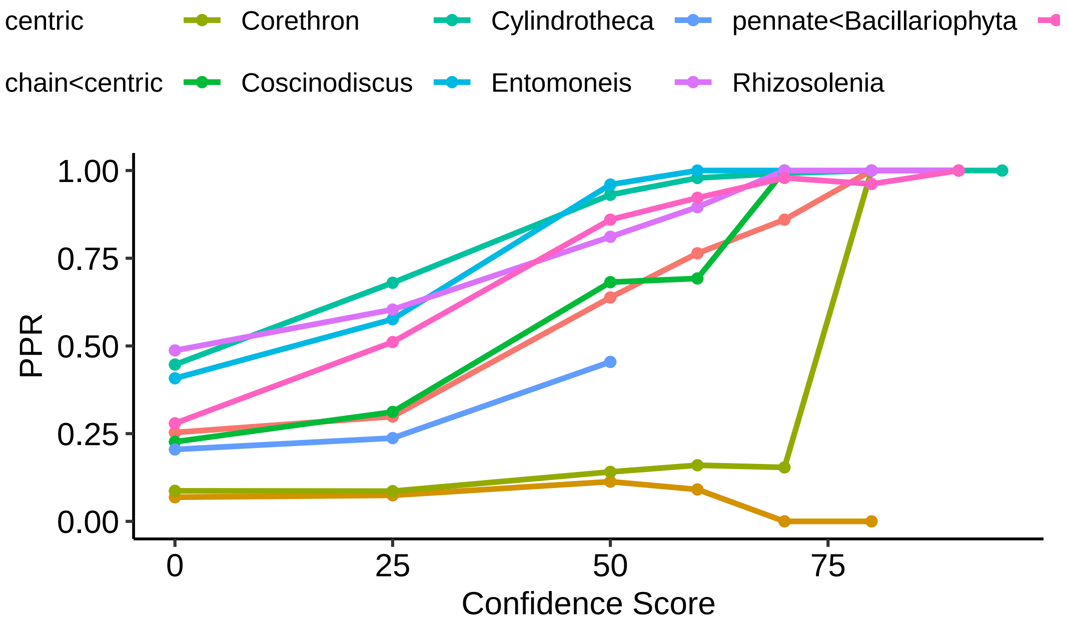


Figure 1: Positive Predictive Rate for different taxonomic classes across a range of confidence scores from Ecotaxa output for the 10x lens. Only taxa which had over 100 predictions were considered.

An acceptable PPR was set at 0.8 so the minimum threshold was identified for 7 taxonmic groups (Table 1).

Table 1: Threshold analysis results for taxonomic groups identified in the 10x FlowCam lens. Shown is the minimum confidence score threshold at which the PPR exceeded 0.8.

| PPR | Score | Taxa-Group |
|---|---|---|
| 0.8600000 | 70 | centric |
| 1.0000000 | 80 | Corethron |
| 1.0000000 | 70 | Coscinodiscus |
| 0.9308462 | 50 | Cylindrotheca |
| 0.9600000 | 50 | Entomoneis |
| 0.8110749 | 50 | Rhizosolenia |
| 0.8600000 | 50 | Skeletonema |

###### 4x Precision.

Gerenally, 4x predicion was poorer. This was in part because the learning set for the size-filtered dataset was much smaller. As a consequence, fewer predictions were made, making the calculation of PPR more challenging. Only considering the taxa with at least 10 predictions in the evaluation set, there were 6 groups. While many of these achieved a high PPR at low confidence scores (0.5), there was a markable drop in PPR once higher scores were considered (Figure 2). Therefore, the only taxa which was reliably identified was *Rhizosolenia spp.* However, prediction of these cells had a PPR greater than 0.8 across all confidence scores.

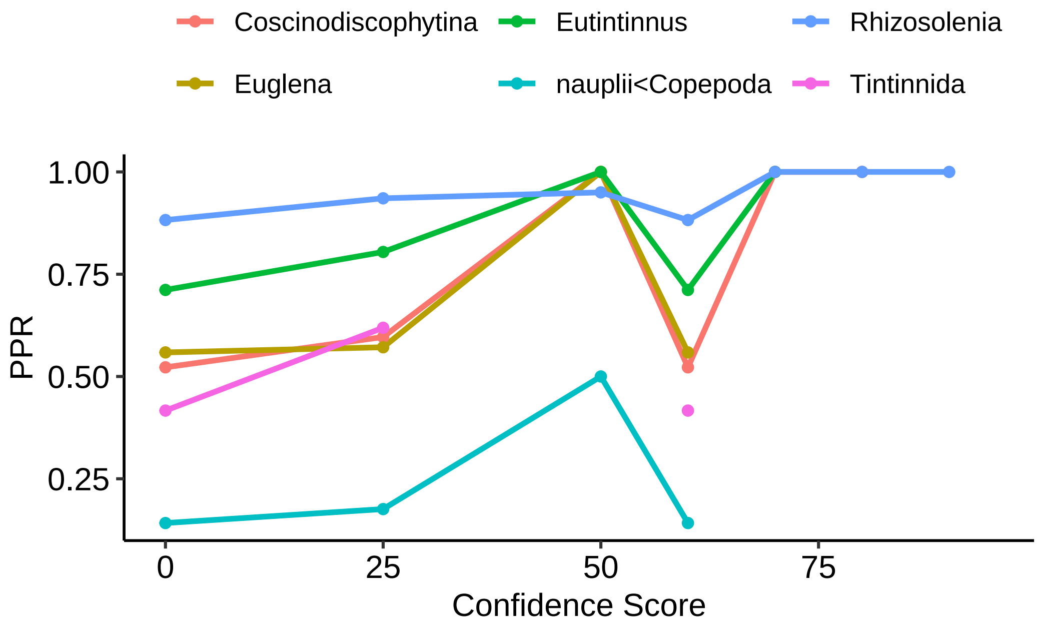


Figure 2: Positive Predictive Rate for different taxonomic classes across a range of confidence scores from Ecotaxa output for the 4x lens. Only taxa which have more than 10 predictions were considered.

Table 2: Positive Predictive Rate for taxa at the 4x lens. Shown is the minimum confidence at which PPR surpassed 0.8. It should be noted however, that for all groups asides from Rhizolenia, the PPR declined below 0.8 at some points when the score increased and more images were included.

| PPR | Score | Taxa-Group |
|---|---|---|
| 1.0000000 | 50 | Coscinodiscophytina |
| 1.0000000 | 50 | Euglena |
| 0.8043478 | 25 | Eutintinnus |
| 0.8823529 | 0 | Rhizosolenia |

#### Taxonomic References:

Horner, R.A. 2002. A Taxonomic Guide to Some Common Marine Phytoplankton. 195 pp. Biopress Ltd., Bristol, England

Larink O. Westheide W. 2011 *Coastal Plankton. Photo Guide for European Seas.* 2nd Ed. Verlag Dr. Friedrich Pfeil, Munchen Germany.

Tomas, C.R. (Editor). 1997. Identifying Marine Phytoplankton. 858 pp. Academic Press, Inc., San Diego, California, USA

https://phytoplanktonguide.lumcon.edu https://www.inaturalist.org/guides/1633
