## Supporting Information 3 for "Storm in a bottle: An experimental investigation of how extreme precipitation impacts phytoplankton communities"

### Posterior Tables

Table 1: Posterior distribution mean and 95% credible interval bounds for different HPLC final mean estimates. Treatments are: A=Control, B=Dilution, C=Salinity, D=Nutrient, E=Composition

| Group | treatment | mean | low.95 | high.95 |
|---|---|---|---|---|
| total_chl | A | 4.5645217 | 2.9897244 | 6.1224819 |
| total_chl | B | 4.2540920 | 2.7333004 | 5.7913066 |
| total_chl | C | 3.5335617 | 1.9672148 | 5.1390079 |
| total_chl | D | 10.1977850 | 8.3543871 | 11.7598770 |
| total_chl | E | 11.0180532 | 9.0754031 | 12.5492905 |
| Chlorophytes | A | 0.5693193 | 0.3772714 | 0.7607573 |
| Chlorophytes | B | 0.5006067 | 0.3107675 | 0.6927519 |
| Chlorophytes | C | 0.5809019 | 0.3897739 | 0.7780218 |
| Chlorophytes | D | 2.2342818 | 2.0441148 | 2.4270699 |
| Chlorophytes | E | 2.3926070 | 2.1972110 | 2.5855403 |
| Cryptophytes | A | 0.1720026 | 0.1205076 | 0.2241694 |
| Cryptophytes | B | 0.1531132 | 0.1006252 | 0.2048727 |
| Cryptophytes | C | 0.1415420 | 0.0890619 | 0.1945694 |
| Cryptophytes | D | 0.3080775 | 0.2565905 | 0.3594633 |
| Cryptophytes | E | 0.4240592 | 0.3714977 | 0.4759215 |
| Cyanobacteria | A | 0.2285451 | 0.1731820 | 0.2846723 |
| Cyanobacteria | B | 0.1402960 | 0.0846131 | 0.1969225 |
| Cyanobacteria | C | 0.2860493 | 0.2291195 | 0.3437774 |
| Cyanobacteria | D | 0.4282072 | 0.3706429 | 0.4877109 |
| Cyanobacteria | E | 0.7365254 | 0.6788494 | 0.7921632 |
| Diatoms | A | 3.5712646 | 2.3161049 | 4.8662558 |
| Diatoms | B | 3.4016297 | 2.1396925 | 4.7330518 |
| Diatoms | C | 2.5286270 | 1.2406544 | 3.9031283 |
| Diatoms | D | 7.7104858 | 6.3403130 | 8.9705187 |
| Diatoms | E | 8.0114953 | 6.6153388 | 9.2771379 |
| Dinoflagellates | A | 0.0219444 | -0.0594555 | 0.1084073 |

| Group | treatment | mean | low.95 | high.95 |
|---|---|---|---|---|
| Dinoflagellates | B | 0.0500600 | -0.0300008 | 0.1326238 |
| Dinoflagellates | C | 0.0496586 | -0.0311095 | 0.1315989 |
| Dinoflagellates | D | 0.0473341 | -0.0347191 | 0.1298469 |
| Dinoflagellates | E | 0.1341581 | 0.0526723 | 0.2134152 |
| Prasinophytes | A | 0.0862045 | 0.0605063 | 0.1123792 |
| Prasinophytes | B | 0.0606947 | 0.0350591 | 0.0863861 |
| Prasinophytes | C | 0.0087420 | -0.0169883 | 0.0339609 |
| Prasinophytes | D | 0.0005089 | -0.0258747 | 0.0263221 |
| Prasinophytes | E | 0.0070242 | -0.0191016 | 0.0337309 |
| Euglenophytes | A | 0.0105128 | -0.0016144 | 0.0222259 |
| Euglenophytes | B | 0.0269544 | 0.0147561 | 0.0391606 |
| Euglenophytes | C | 0.0060425 | -0.0056206 | 0.0178003 |
| Euglenophytes | D | 0.0037374 | -0.0081811 | 0.0158153 |
| Euglenophytes | E | 0.0004737 | -0.0111089 | 0.0121570 |
| Haptophytes | A | 0.0326244 | 0.0205193 | 0.0449675 |
| Haptophytes | B | 0.0229570 | 0.0108843 | 0.0346667 |
| Haptophytes | C | 0.0078476 | -0.0047702 | 0.0200931 |
| Haptophytes | D | 0.1036713 | 0.0915471 | 0.1163925 |
| Haptophytes | E | 0.0418997 | 0.0296876 | 0.0540204 |

Table 2: Posterior distribution mean and 95% credible interval bounds for different HPLC final chlorophyll-a change estimates. Treatments are: A=Control, B=Dilution, C=Salinity, D=Nutrient, E=Composition

| Group | treatment | mean | low.95 | high.95 |
|---|---|---|---|---|
| total_chl | A | -2.2421395 | -3.8072204 | -0.7088135 |
| total_chl | B | 0.1402345 | -1.4300766 | 1.6844470 |
| total_chl | C | 0.1897291 | -1.4013310 | 1.7978130 |
| total_chl | D | 8.2078652 | 6.5993667 | 9.8016116 |
| total_chl | E | 6.3981016 | 4.7879219 | 7.9599279 |
| Chlorophytes | A | -0.2444002 | -0.4331016 | -0.0617346 |
| Chlorophytes | B | 0.0869578 | -0.1008345 | 0.2742696 |
| Chlorophytes | C | 0.2085090 | 0.0204696 | 0.4049070 |
| Chlorophytes | D | 1.9000907 | 1.7084825 | 2.0892915 |
| Chlorophytes | E | 1.5775933 | 1.3862051 | 1.7738311 |
| Cryptophytes | A | -0.0002847 | -0.0528495 | 0.0525046 |
| Cryptophytes | B | -0.0173190 | -0.0706879 | 0.0338743 |
| Cryptophytes | C | 0.0242405 | -0.0267446 | 0.0753742 |
| Cryptophytes | D | 0.1759439 | 0.1238647 | 0.2264798 |
| Cryptophytes | E | 0.1161837 | 0.0647706 | 0.1665758 |

| Group | treatment | mean | low.95 | high.95 |
|---|---|---|---|---|
| Cyanobacteria | A | 0.0606129 | 0.0052150 | 0.1151942 |
| Cyanobacteria | B | 0.0319796 | -0.0257451 | 0.0874781 |
| Cyanobacteria | C | 0.2043651 | 0.1468289 | 0.2606069 |
| Cyanobacteria | D | 0.3473846 | 0.2905551 | 0.4048038 |
| Cyanobacteria | E | 0.5854220 | 0.5290255 | 0.6417157 |
| Diatoms | A | -1.4092713 | -2.7737634 | -0.0395420 |
| Diatoms | B | 0.6951764 | -0.6790492 | 2.0410205 |
| Diatoms | C | 0.4083145 | -0.9454229 | 1.7850729 |
| Diatoms | D | 6.3323521 | 4.9637149 | 7.7050354 |
| Diatoms | E | 5.0363595 | 3.7055025 | 6.3804095 |
| Dinoflagellates | A | -0.0209394 | -0.1034867 | 0.0611264 |
| Dinoflagellates | B | -0.0426998 | -0.1241141 | 0.0409566 |
| Dinoflagellates | C | -0.0608610 | -0.1452979 | 0.0224985 |
| Dinoflagellates | D | -0.0725123 | -0.1570705 | 0.0106778 |
| Dinoflagellates | E | -0.1798928 | -0.2631509 | -0.0946284 |
| Prasinophytes | A | -0.2280251 | -0.2540666 | -0.2024278 |
| Prasinophytes | B | -0.1211818 | -0.1462241 | -0.0951823 |
| Prasinophytes | C | -0.1340831 | -0.1601914 | -0.1080075 |
| Prasinophytes | D | -0.1436759 | -0.1696549 | -0.1180317 |
| Prasinophytes | E | -0.3515162 | -0.3776487 | -0.3254580 |
| Euglenophytes | A | -0.3955970 | -0.4076851 | -0.3832705 |
| Euglenophytes | B | -0.5329532 | -0.5452321 | -0.5208192 |
| Euglenophytes | C | -0.4540624 | -0.4657420 | -0.4420861 |
| Euglenophytes | D | -0.3622202 | -0.3741500 | -0.3505910 |
| Euglenophytes | E | -0.3765932 | -0.3883608 | -0.3648740 |
| Haptophytes | A | -0.0423838 | -0.0545581 | -0.0306902 |
| Haptophytes | B | 0.0156500 | 0.0038017 | 0.0276062 |
| Haptophytes | C | -0.0001664 | -0.0126058 | 0.0116390 |
| Haptophytes | D | 0.1033358 | 0.0910876 | 0.1156226 |
| Haptophytes | E | 0.0399442 | 0.0279091 | 0.0520348 |

Table 3: Posterior distribution mean and 95% credible interval bounds for different diatom carbon biomass concentration final mean estimates. Treatments are: A=Control, B=Dilution, C=Salinity, D=Nutrient, E=Composition

| Group | treatment | mean | low.95 | high.95 |
|---|---|---|---|---|
| Rhizosolenia | A | 1.796460e+07 | 10128762.6449 | 23923030.055 |
| Rhizosolenia | B | 1.345411e+07 | 6659764.0719 | 18729337.447 |
| Rhizosolenia | C | 3.509031e+06 | -1285562.2993 | 8088076.079 |
| Rhizosolenia | D | 5.778880e+06 | 674991.0280 | 10418120.554 |

| Group | treatment | mean | low.95 | high.95 |
|---|---|---|---|---|
| Rhizosolenia | E | 6.189462e+06 | 1059906.8516 | 10819479.870 |
| Skeletonema | A | 1.695736e+06 | 1118002.6693 | 2275948.464 |
| Skeletonema | B | 2.802085e+06 | 2228431.7142 | 3398546.129 |
| Skeletonema | C | 4.911648e+05 | -93076.9014 | 1078248.112 |
| Skeletonema | D | 1.153287e+06 | 575580.3138 | 1743529.694 |
| Skeletonema | E | 2.304306e+06 | 1682848.6488 | 2921938.282 |
| Entomoneis | A | 4.213513e+05 | -1616452.1257 | 2439723.464 |
| Entomoneis | B | 2.938498e+05 | -1727824.7229 | 2435855.086 |
| Entomoneis | C | 2.905236e+05 | -1685303.2895 | 2382869.153 |
| Entomoneis | D | 3.730778e+05 | -1609292.4204 | 2401765.019 |
| Entomoneis | E | 8.388285e+06 | 6276854.7222 | 10394861.400 |
| Cylindrotheca | A | 4.328381e+05 | -1649979.6688 | 2564488.301 |
| Cylindrotheca | B | 5.943895e+05 | -1505189.2109 | 2701802.888 |
| Cylindrotheca | C | 3.467965e+05 | -1652516.9813 | 2415342.397 |
| Cylindrotheca | D | 4.474951e+05 | -1605094.3991 | 2598377.616 |
| Cylindrotheca | E | 1.148436e+07 | 9276823.3929 | 13453987.580 |
| centric | A | 8.838163e+05 | 534888.8633 | 1225358.735 |
| centric | B | 2.329835e+05 | -111272.5480 | 578218.283 |
| centric | C | 3.927046e+05 | 52546.9504 | 736827.047 |
| centric | D | 7.876195e+05 | 444126.8852 | 1135916.297 |
| centric | E | 5.061951e+05 | 158843.3463 | 847789.297 |
| Coscinodiscus | A | 8.278901e+06 | 5118114.9972 | 11265464.337 |
| Coscinodiscus | B | 1.528075e+06 | -1454960.5214 | 4581537.510 |
| Coscinodiscus | C | 1.558103e+06 | -1371802.5854 | 4529008.946 |
| Coscinodiscus | D | 1.271608e+06 | -1704680.5314 | 4271671.913 |
| Coscinodiscus | E | 4.102942e+06 | 1001021.4258 | 7162021.337 |
| Corethron | A | 4.191698e+03 | 2180.0466 | 6237.261 |
| Corethron | B | 2.443469e+03 | 379.1922 | 4451.618 |
| Corethron | C | 4.562932e+02 | -1512.0252 | 2479.037 |
| Corethron | D | 1.033395e+03 | -901.2349 | 3045.503 |
| Corethron | E | -4.275720e+00 | -2072.9433 | 2104.534 |

Table 4: Posterior distribution mean and 95% credible interval bounds for different diatom carbon biomass concentration change ( $pgCmL^{-1}$ ) estimates. Treatments are: A=Control, B=Dilution, C=Salinity, D=Nutrient, E=Composition

| Group | treatment | mean | low.95 | high.95 |
|---|---|---|---|---|
| Rhizosolenia | A | 8.983089e+06 | -10318055.156 | 28398103.27 |
| Rhizosolenia | B | 1.036986e+07 | -9368070.757 | 29790041.13 |
| Rhizosolenia | C | 1.023211e+06 | -18053342.235 | 19943714.77 |

| Group | treatment | mean | low.95 | high.95 |
|---|---|---|---|---|
| Rhizosolenia | D | 1.894454e+06 | -17401478.691 | 20650168.15 |
| Rhizosolenia | E | 1.976278e+06 | -17470070.458 | 20357612.62 |
| Skeletonema | A | 1.234449e+07 | 6827384.863 | 17628667.41 |
| Skeletonema | B | 2.209106e+07 | 16216562.603 | 27456443.66 |
| Skeletonema | C | 3.128814e+06 | -2177390.432 | 8451716.28 |
| Skeletonema | D | 9.470618e+06 | 4060271.395 | 14849750.45 |
| Skeletonema | E | 2.008087e+07 | 14444122.398 | 25321806.09 |
| Entomoneis | A | 1.048260e+06 | -14379628.169 | 16537212.63 |
| Entomoneis | B | 1.099805e+06 | -14871754.180 | 17082428.66 |
| Entomoneis | C | 7.993733e+05 | -14879619.422 | 16616851.10 |
| Entomoneis | D | 1.222029e+06 | -14704791.126 | 16544067.18 |
| Entomoneis | E | 2.796943e+07 | 7788484.134 | 47756949.53 |
| Cylindrotheca | A | 1.200476e+06 | -14296638.807 | 16654803.91 |
| Cylindrotheca | B | 1.822349e+06 | -13436242.973 | 17257325.14 |
| Cylindrotheca | C | 8.726624e+05 | -14435028.251 | 16026163.04 |
| Cylindrotheca | D | 1.327482e+06 | -13829700.661 | 16342966.63 |
| Cylindrotheca | E | 3.254235e+07 | 10382573.564 | 53702412.62 |
| centric | A | 4.311450e+06 | 1418217.500 | 7237300.14 |
| centric | B | 1.392910e+05 | -2670526.005 | 3027588.53 |
| centric | C | 1.974741e+06 | -774051.808 | 4787979.87 |
| centric | D | 4.373875e+06 | 1454044.636 | 7180272.27 |
| centric | E | 2.253806e+06 | -496651.911 | 5008194.95 |
| Coscinodiscus | A | 1.344188e+07 | -6516891.499 | 33039950.24 |
| Coscinodiscus | B | 2.732534e+06 | -15953913.287 | 21370033.91 |
| Coscinodiscus | C | 2.645412e+06 | -16001588.929 | 21502404.00 |
| Coscinodiscus | D | 2.159462e+06 | -16589274.531 | 20751532.21 |
| Coscinodiscus | E | 6.868796e+06 | -11794608.158 | 24984563.77 |
| Corethron | A | 2.643817e+04 | 13432.652 | 39004.88 |
| Corethron | B | 1.496403e+04 | 2294.518 | 27462.93 |
| Corethron | C | 2.554364e+03 | -10526.006 | 15907.75 |
| Corethron | D | 6.705364e+03 | -5972.290 | 19481.30 |
| Corethron | E | 5.949349e+01 | -12381.041 | 12694.40 |
